# Osteoarthritis Early-, Mid- and Late-Stage Progression in the Rat Medial Meniscus Transection Model

**DOI:** 10.1101/2021.03.11.434909

**Authors:** Thanh N. Doan, Jay M. McKinney, Krishna A Pucha, Fabrice C. Bernard, Nick J. Willett

**Author notes:** **Corresponding Author:** Nick J. Willett, **Email:**, **Address:** 1670 Clairmont Road, Room 5A-115, Decatur, GA 30033. Thanh Doan and Jay McKinney have contributed equally to the work and shared first authorship designation. **Emails for Authors** Thanh Doan, Jay McKinney, Krishna Pucha, Fabrice Bernard, Nick Willett.

## Abstract

Osteoarthritis is a degenerative disease of synovial joints affecting all tissues, including the articular cartilage and underlying subchondral bone. Osteoarthritis animal models can recapitulate aspects of human disease progression and are commonly used to test the development of drugs, biomaterials, and cell therapies for treatment. The rat medial meniscus transection (MMT) model is a surgically induced post-traumatic osteoarthritis model and is one of the most commonly used models for therapeutic development; however, it is typically used to evaluate the efficacy of therapies to prevent disease development rather than testing the treatment of disease progression in already established disease. We describe herein, the qualitative and quantitative changes to articular cartilage, subchondral bone, and formation of osteophytes in rats at early-(3-weeks post-surgery), mid-(6-weeks post-surgery) and late-(12-weeks post-surgery) stages of osteoarthritis progression. Tibiae of MMT-operated animals showed loss of proteoglycan and fibrillation formation on articular cartilage surfaces as early as 3-weeks post-surgery. Using a contrast-enhanced μCT technique, quantitative, 3-dimensional analysis of the tibiae showed that the articular cartilage initially thickened at 3- and 6-weeks post-surgery and then decreased at 12-weeks post-surgery. This decrease in cartilage thickness corresponded with increased lesions in the articular cartilage, including fully degraded surfaces down to the subchondral bone layer. In this rat MMT model, subchondral bone thickening was significant at 6-weeks post-surgery and seem to follow cartilage damage. Osteophytes were found at 3-weeks post-surgery, which coincided with articular cartilage degradation. Cartilaginous osteophytes preceded mineralization suggesting that these marginal tissue growths most likely occurred through endochondral ossification. The use of the rat MMT model has predominantly been used out to 3-weeks, and most studies determine the effect of therapies to delay or prevent the onset of osteoarthritis. We provide evidence that an extension of the rat MMT model out to 6 and 12 weeks resembled more severe phenotypes of human osteoarthritis. The mid- to late-stages of rat MMT model can be used to evaluate the therapeutic efficacy of novel treatments to treat the progression of established disease — since patients typically present in the clinic when the disease is established and becomes symptomatic, thus evaluating the efficacy of new treatments at the late stage will be important for eventual clinical translation.

## Introduction

Osteoarthritis is a debilitating disease of joints that presents with progressive changes to synovium, articular cartilage degradation, and subchondral bone remodeling. Patients with osteoarthritis will most commonly seek treatment from a physician because of joint pain [1], joint effusion and/or inability to move freely. Physicians rely on joint pain assessment [2, 3], radiography, ultrasound [4, 5], and other imaging modalities to diagnose osteoarthritis [6]. The most common method employed is x-ray to visualize joint bones and the joint space [7]; however, this technique does not directly assess the cartilage, which is one of the first tissues that degenerate during early osteoarthritis. The diagnosis of osteoarthritis in the clinic, on the other hand, is usually at a more advanced stage of the disease, where narrowing of the joint space is already present on x-ray images, and bone spurs are sometimes already visible. By the time a patient presents in the clinic, the clinician will be treating established disease and trying to mitigate the symptoms.

For research and development of therapeutics to treat osteoarthritis, numerous animal models [8] have been used to mimic and model the degenerative disease [9]. These pre-clinical models have been used to understand the biological development of osteoarthritis and to test putative therapeutics [10]. Unfortunately, to date, no disease modifying therapeutic is available, and this may be, in part, due to a lack of predictive capacity and utility of osteoarthritis animal models. The induction of osteoarthritis in animal models is typically achieved by one of three primary means: 1) spontaneously, 2) chemically, or 3) mechanically. Spontaneous osteoarthritis either relies on animals that naturally get the disease (e.g., guinea pigs and horses) or relies on manipulating the genetic background of the animal and/or aging of the animal; spontaneous osteoarthritis tend to take a long time (e.g. months to years) for onset of the disease and models primary osteoarthritis development. Chemical induction of osteoarthritis relies on local application of catabolic agent(s) to quickly (e.g., days or weeks) cause degeneration of the joint. Even though chemical induction may induce some of the key osteoarthritis phenotypes, such as synovitis, degenerated cartilage, pain, and bone remodeling, the etiology and progression of the disease tend to be very different than human osteoarthritis. So chemical-induced models are less commonly utilized for testing therapeutics that target disease development and more for therapeutics that focuses on treating symptoms, e.g., pain. Further, overdose or overexposure of catabolic agent(s) may cause a strong inflammatory response in the affected joint and may result in a disease more similar to rheumatoid arthritis than osteoarthritis. The mechanical method typically entails a surgical manipulation to destabilize the joint, which leads to post-traumatic osteoarthritis. The amount of time (typically weeks to months) for osteoarthritic features to develop depends on the type and size of the animal and the severity of joint destabilization (i.e., partial or full transection of the meniscus or multiple ligaments being transected). Post-traumatic osteoarthritis animal models tend to develop osteoarthritis faster than most genetic spontaneous animal models and have biological relevance to secondary osteoarthritis in humans. The rat medial meniscus transection (MMT) osteoarthritis model is one of the most common surgically-induced models and is often utilized to phenotypically characterize the early stages of osteoarthritis development [11, 12] and to test therapeutics [13, 14]. However, the majority of work with this model is at the early stages of the disease and testing therapeutics used to prevent disease development or onset rather than evaluating or testing the progression or treatment of established disease.

Potential therapeutics, including small molecules [15, 16], biomaterials [17, 18], and cell-based administration [13, 19, 20], have been tested in rats; however, most of the regimens were given days immediately after the surgical procedure which would delay or prevent the onset of the disease. Very few studies have investigated the efficacy of therapeutics after osteoarthritis has already developed within the joint [21], and even then, the time of therapeutic intervention was not related to human clinical manifestation. The testing of these therapeutics at the onset or early stage of osteoarthritis development in animal models may not translate to clinical benefit and thus potentially contributing to the fact that no disease-modifying drug has been identified for osteoarthritis. The characterization of the disease progression in this model has largely been at early timepoints and stages; there is not a well-defined and comprehensive understanding of the progression to mid- and late-stage disease with this model. Our objective was to quantitatively evaluate osteoarthritis progression in the rat MMT model from the commonly utilized early-stage disease time point to mid- and late-stage disease. With more recent advances with small animal imaging modalities, like contrast enhanced μCT, we have provided herein quantitative analysis of morphological changes to the articular cartilage and subchondral bone during differing stages of post-traumatic osteoarthritis. By defining the stages of osteoarthritis progression in the rat MMT osteoarthritis model, the use of therapies to treat the disease may be better realized and more closely resemble the clinic manifestation.

## Materials and Methods

### Surgical Methods

All procedures followed institutional guidelines set by the Atlanta Veterans Affairs Medical Center (VAMC) and were approved by Institutional Animal Care and Use Committee (IACUC). Twelve week old male Lewis rats (Charles River) were acclimated for one-week post-arrival and surgical manipulation occurred the following week.

The medial meniscal transection (MMT) is a surgical procedure to destabilize the rat knee joint and replicates the pathology of osteoarthritis (Figure 1) [8]. Briefly, animals were anesthetized via isoflurane inhalation, and the surgical site was shaved and sterilized immediately prior to surgery. Sustained-release (SR) buprenorphine (0.03 mg/kg) (ZooPharm, Windsor, CO) was injected subcutaneously as an analgesic. Skin and soft tissue incision were made along the sterile surgical site of the medial left hindlimb, along the femora-tibial joint. A blunt dissection exposed the medial collateral ligament (MCL), which was transected to expose the meniscus. For MMT animals, the meniscus was fully cut through at the narrowest point. For Sham animals, only the MCL was transected, and the meniscus was left intact. Soft tissues were closed with 4.0 vicryl sutures, and the skin was closed with staple clips. The contralateral right hindlimbs were unoperated and served as naïve controls herein. Post-surgery, animals were rehydrated with 10 mL of Lactated Ringer’s solution. All animals were monitored for three consecutive days post-surgery. Staples were removed one week after surgery when the skin had fully closed and healed.

**Figure 1:**
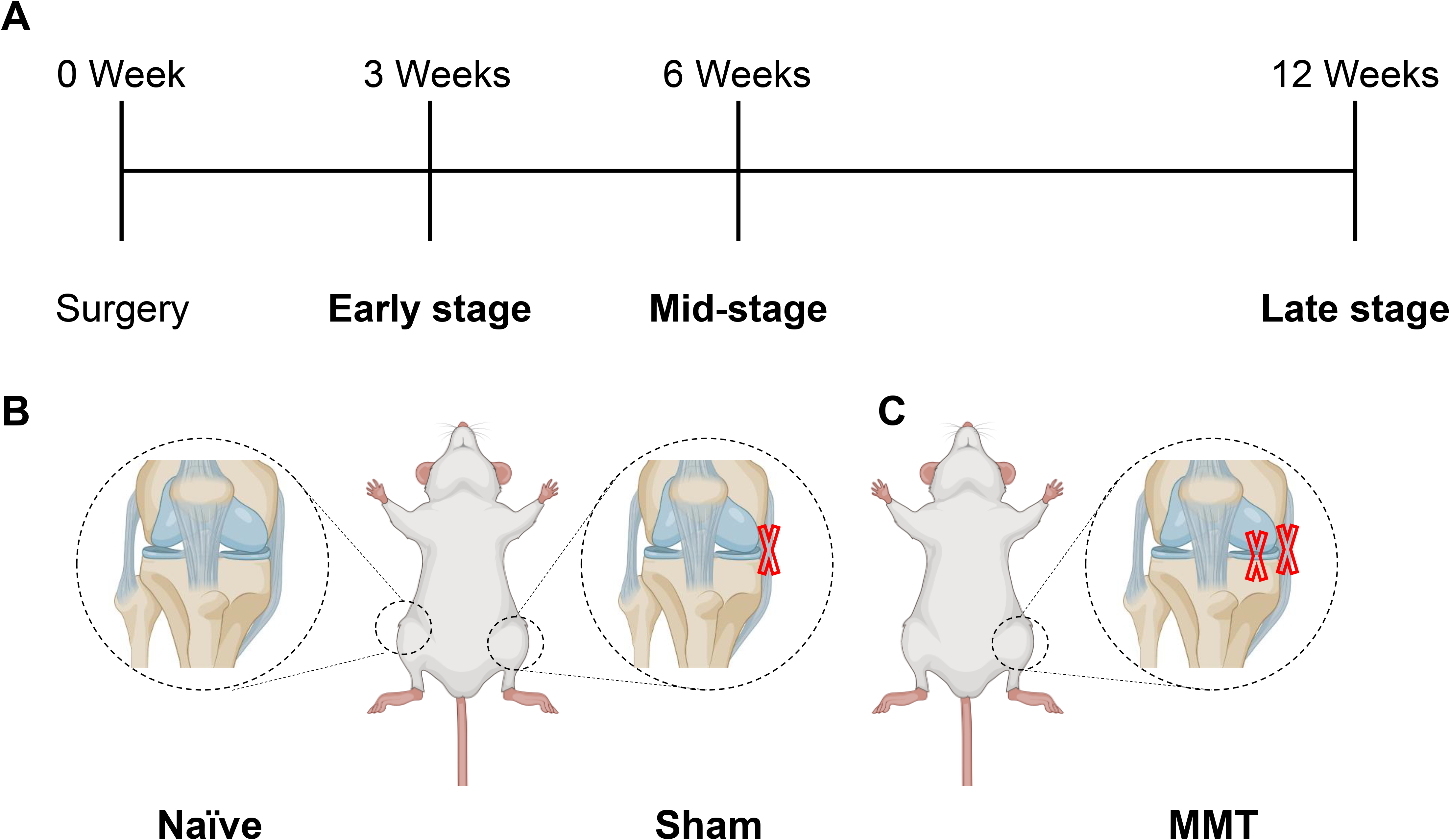
Illustration of Naïve, Sham- and MMT-operated hindlimbs in rats. Lewis rats were anaesthetized with isoflurane and their hindlimbs either received no surgery (naïve, right hindlimb), Sham-operation (transection of medial collateral ligament on left hindlimb), or MMT-operation (transection of medial collateral ligament and medial meniscus on left hindlimb). After surgical manipulation, soft tissues were closed with 4.0 vicryl suture and the skin were closed with staples. Animals recovered from anesthesia and were returned to their housing. Rats were euthanized at 3-, 6- and 12-weeks post-surgery to determine the extent of osteoarthritis damage.

### Preparation of hindlimbs for microCT and for histology

Animals were euthanized at different timepoints post-surgery (3-, 6- and 12-weeks) via CO_2_ asphyxiation. Cervical dislocation was used as a secondary euthanasia method after asphyxiation. Left and right hindlimbs were dissected and fixed in 10% neutral buffered formalin. Muscle and connective tissues were removed from the hindlimbs. The femur was disarticulated from the tibia. Meniscus and residual soft tissue surrounding the medial tibial condyle were dissected and discarded.

### EPIC-μCT analysis of articular cartilage, osteophytes, and subchondral bone

Equilibrium partitioning of an ionic contrast agent-based micro-computed tomography (EPIC-μCT) was used to quantitatively assess structural and compositional parameters of articular cartilage, subchondral bone, and osteophytes of tibiae. Tibiae were immersed in 30% Hexabrix 320 contrast reagent (NDC 67684-5505-5, Guerbet, Villepinte, France) for 30 min at 37°C before being scanned on Scanco μCT 40 (Scanco Medical, Brüttisellen, Switzerland). The following parameters were used for acquiring images of proximal tibiae: 45 kVp, 177 μA, 200 ms integration time and isotropic 16 μm voxel size.

Two-dimensional (2D) grayscale tomograms were reconstructed from raw data and were orthogonally transposed to create a 3D reconstruction of the joint (Figure 2A). 2D coronal sections were used to contour the volume of interest (VOI) of each joint. The VOI for contoured articular cartilage (Figure 2C and 2D), subchondral bone (Figure 2F and 2G) and osteophytes (Figure 2I and 2J) were delineated from the background and soft tissues by setting lower and upper thresholds. Contoured regions were evaluated for volume, thickness, and attenuation along the full length and medial 1/3 region of articular cartilage and subchondral bone. The medial 1/3 region exhibited the greatest change to articular cartilage and subchondral bone from MMT surgery. Articular cartilage attenuation was shown to be inversely proportional to sulfated glycosaminoglycan (sGAG) content, a key component of the cartilage matrix [22]. Increased partitioning of Hexabrix 320 reagent resulted in increased attenuation within the damaged cartilage region. Osteophyte volumes were measured as previously described [11, 13].

**Figure 2:**
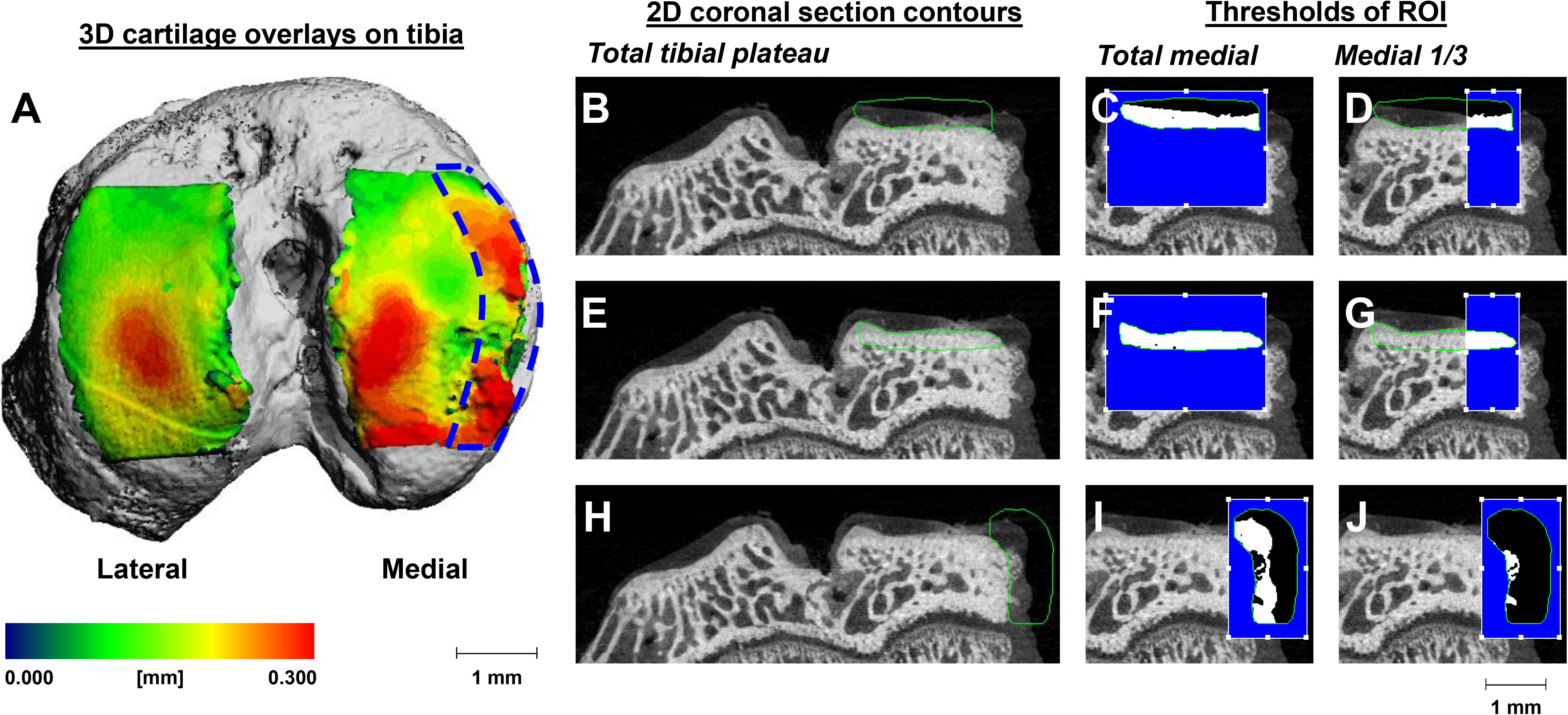
Representation of EPIC-microCT analysis of articular cartilage and osteophytes. A) Proximal view of a rat tibia isolated from an MMT-operated hindlimb at 3-weeks post-surgery. Lateral condyle exhibited even smooth surface of the articular cartilage while the medial condyle showed irregularity of articular cartilage and several regions of lesions. The blue dashed area marked the medial 1/3 region of the medial condyle. B) Representative image of a 2D coronal-sectioned EPIC-μCT image of a tibia isolated from a 3-weeks MMT-operated hindlimb. Articular cartilage of the medial condyle was contoured (green outline). The total medial articular cartilage was analyzed (C) and the medial 1/3 region of the articular cartilage was analyzed (D). The white region within the green contoured area represented the analyzed area that were within the set threshold and the black region within the green contoured area represented the area not analyzed and were outside the set threshold. E) Same image as panel B with green contour outlining the subchondral bone and F) the total subchondral bone region for analysis and G) the medial 1/3 subchondral bone region. H) Same image as panel B with green contour outlining the osteophyte and I) the cartilaginous osteophyte region and J) the mineralized osteophyte region.

### Surface roughness analysis of articular cartilage

Serial 2D images of the proximal tibiae were analyzed using a customized algorithm in MATLAB (MathWorks, Natick, MA) to quantify surface roughness, lesion volume, and full-thickness lesion area [23]. Images were processed to generate a 3D surface of the articular cartilage surface (Figure 3). This 3D rendering was fitted along a 3D polynomial surface: fourth-order along the ventral-dorsal axis and second-order along the medial-lateral axis [23]. The root mean square difference between the generated (actual) and polynomial fitted (predicted) surfaces was the measure of cartilage layer surface roughness (Figure 3D-G). Lesion volume (Figure 3H-K) was calculated as the volume of root mean square difference between the generated fitted surface and the polynomial surface where > 25% of total (predicted) cartilage thickness was exceeded. Full-thickness lesion area (Figure 3L-O), also called exposed bone, was the sum of the area on the tibial condyle where no cartilage layer was present. Surface roughness, lesion volume and full-thickness lesion area were calculated for full and medial 1/3 region of the articular cartilage.

**Figure 3:**
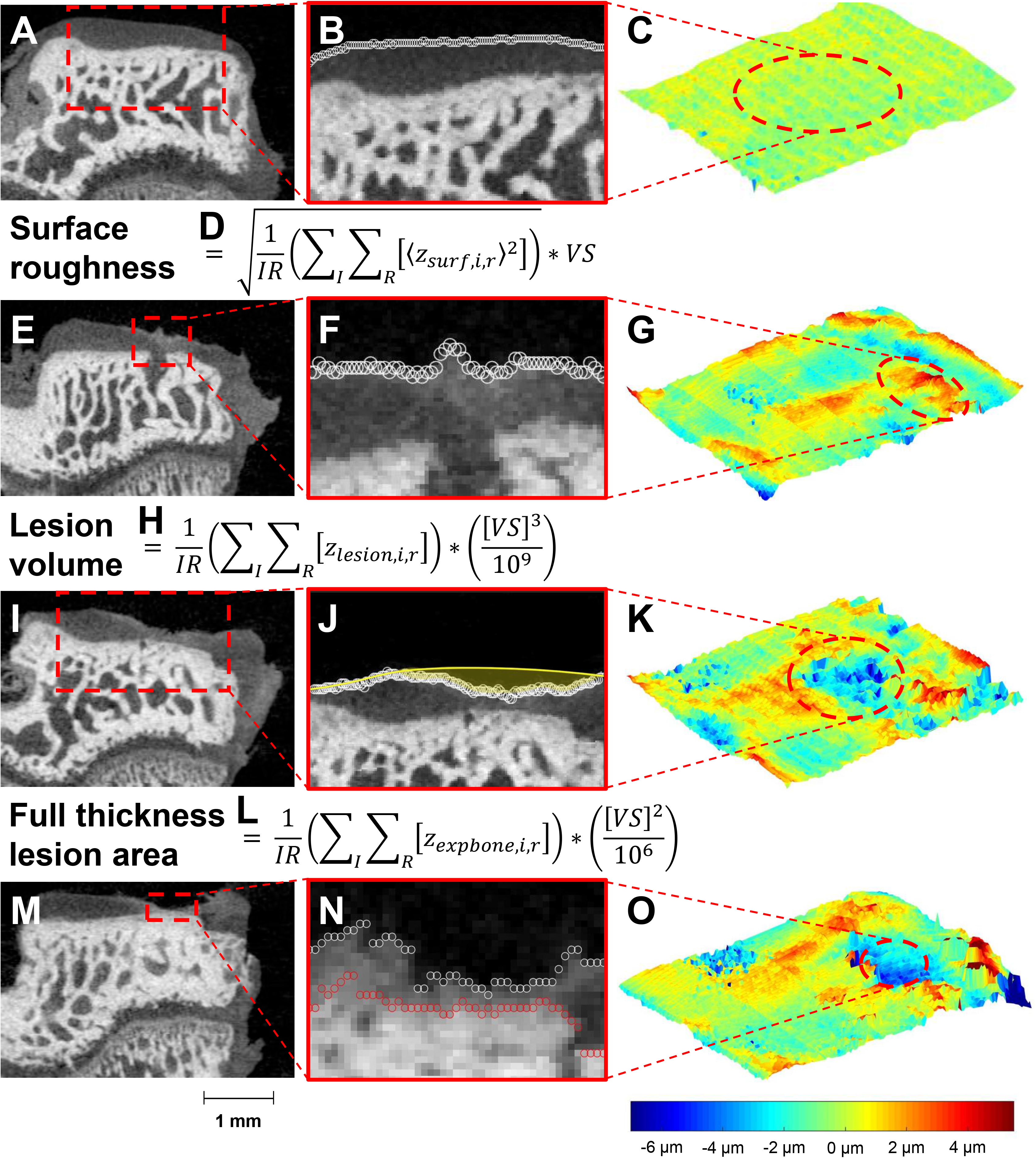
Analysis for surface roughness, lesion volume and full thickness lesion area of articular cartilage. A) Representative image of a 2D coronal-sectioned EPIC-μCT image of a tibia isolated from a 3-weeks Sham-operated hindlimb. The region of the dashed red box was shown at higher magnification (B). The white boundary represented the boundary of the articular cartilage derived from the customized MATLAB code and C) represented the relative deviation from the fit over the proximal plane of the medial condyle (similar to Figure 2A) with the dashed red ellipse highlighting the relative smoothness of the articular cartilage surface. E) Representative image of a 2D coronal-sectioned EPIC-μCT image of a tibia isolated from a 3-weeks MMT-operated hindlimb. The region of the dashed red box was shown at higher magnification (F). The white boundary represented the boundary of the articular cartilage derived from the customized MATLAB code and G) represented the relative deviation from the fit over the proximal plane of the medial condyle with the dashed red ellipse highlighting an uneven surface of the articular cartilage. I) Representative image of a 2D coronal-sectioned EPIC-μCT image of a tibia isolated from a 6-weeks MMT-operated hindlimb. The region of the dashed red box was shown at higher magnification (J). The white boundary represented the boundary of the articular cartilage and the yellow area represented the fit derived from the customized MATLAB code. K) 3D representation of the articular cartilage with the dashed red ellipse highlighting an area with lesions. M) Representative image of a 2D coronal-sectioned EPIC-μCT image of a tibia isolated from a 12-weeks MMT-operated hindlimb. The region of the dashed red box was shown at higher magnification (N). The white boundary represented the boundary of the articular cartilage and its closeness to the subchondral bone (red boundary) derived from the customized MATLAB code. O) 3D representation of the articular cartilage with the dashed red ellipse highlighting a full thickness lesion area or exposed bone. The formulas for surface roughness, lesion volume and full thickness lesion area were shown in D, H and L, respectively.

### Histology

Following μCT analysis, contrast agent was removed from tissues through washes in PBS. Naïve, Sham and MMT tibiae were decalcified with formic/citrate decalcifying solution (Newcomer Supply 10492C, Middleton, WI) for 7 days. Samples were paraffin-embedded and sectioned into 5 μm-thick slices. Sections were stained with hematoxylin and eosin (H&E; Fisherbrand™ 517-28-2, Waltham, MA) or safranin-O and fast green (Saf-O; Electron Microscopy Sciences^®^ 20800, Hatfield, PA), following manufacturer protocols. Representative serial images for H&E stains were included (Supplementary Figure 1).

### Statistical Analysis

All figures were presented as means +/− standard deviation (SD). Two-way analysis of variance (ANOVA) was used to determine significant differences between groups (*p* < 0.05). Tukey Honest post-hoc analysis was used to determine between group differences for cartilage and subchondral bone parameters. Bonferroni non-parametric post-hoc analysis was used for osteophyte parameters. Data were analyzed and plotted using GraphPad Prism software version 6.0 (GraphPad Software Inc., La Jolla, CA).

## Results

### Qualitative and quantitative analysis of articular cartilage

Rats received either Sham or MMT surgery to the left hindlimb knee joint with contralateral right hindlimbs serving as naïve (non-operated) controls. Similar to previous reports for the rat MMT model [13, 14], the articular cartilage of tibia from Sham-operated animals were indistinguishable from naïve hindlimbs at all stages investigated: 3-, 6- and 12-weeks post-surgery (Figure 4), as represented with safranin-O (red) stain for articular cartilage and with quantitative analysis using EPIC-μCT. MMT animals exhibited loss of safranin-O stain in the medial region of the tibia (Figure 4C) at 3-weeks post-surgery. Lack of safranin-O stain indicated a loss of proteoglycan content within the articular cartilage extracellular matrix. In addition, there was evidence of fibrillation on the articular cartilage surface. These changes to the articular cartilage were quantifiable with the EPIC-μCT technique across the full length (Figure 4G-I) and medial 1/3 regions of the cartilage (Figure 4J-L). MMT-operated articular cartilage thickness was significantly increased compared to Sham or naïve tibiae. The increased articular cartilage thickness was most evident in the medial 1/3 region of the tibiae, which was consistent with the joint destabilization by transecting the medial meniscus. Average articular cartilage thickness in the medial 1/3 region decreased at 12-weeks post-surgery for MMT animals due to loss of chondrocytes and extracellular matrix (Figure 6C). Consistent with increased thickness, MMT-operated articular cartilage volume was also increased, presumably due to swelling of the articular cartilage matrix as the proteoglycan matrix was degraded. Surface fibrillations and roughening was quantified by quantitatively evaluating the surface roughness of the articular cartilage (Figure 4I and 4L). Undamaged articular cartilage surfaces were smooth and continuous throughout the tibia condyle, like those found for naïve and Sham animals. The articular cartilage surface from MMT animals showed increased surface roughness due to fibrillation of the surface and the presence of cartilage lesions (Figure 5).

**Figure 4:**
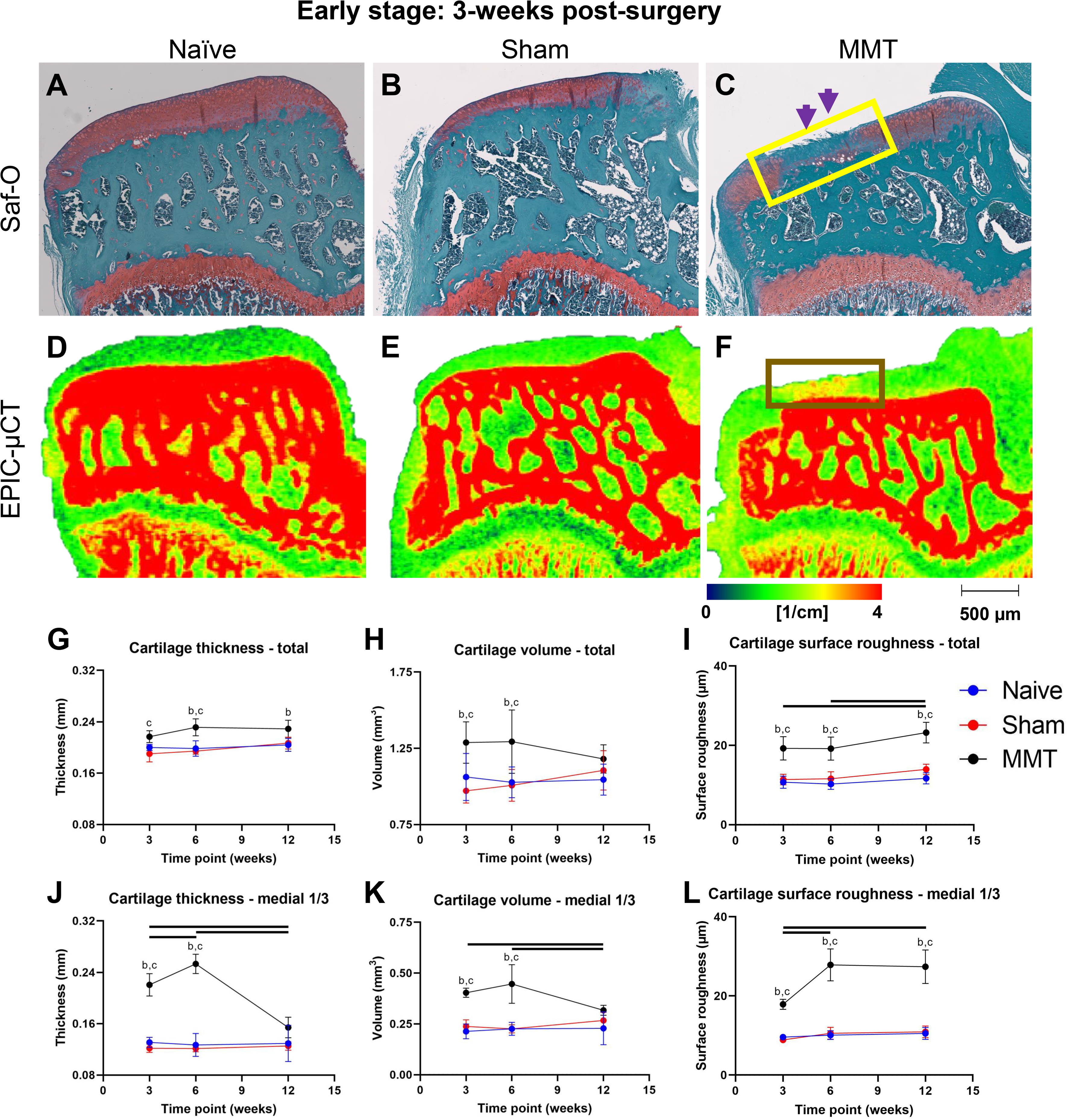
Changes to articular cartilage structure and morphometry during OA progression. Rats were euthanized at 3-weeks following Sham- or MMT-surgery and their hindlimbs were fixed in 4% neutral-buffered formalin. After embedding in paraffin, 5μm coronal serial sections of naïve (A), Sham-operated (B) and MMT-operated (C) tibiae were placed onto glass slides and stained with safranin-O/fast-green. Red safranin-O stain was found in the articular cartilage layer and growth plate. These regions were dominated by chondrocytes and the red color was indicative of proteoglycan matrix. Fast-green was used as a counter stain. The lack of safranin-O stain in the articular cartilage of MMT-operated hindlimbs (yellow box) suggested loss of proteoglycan content. Fibrillation (purple arrow heads) of the matrices was evident and resulted in increased surface roughness of the articular cartilage. 2D images of EPIC-μCT coronal sections that closely matched the safranin-O/fast green histology sections were shown for naïve (D), Sham-operated (E) and MMT-operated (F) tibia. Higher attenuation of the articular cartilage (brown box) was found for tibia of MMT-operated animals. Custom-coded MATLAB algorithms were used to quantify articular cartilage thickness, volume and surface roughness across the total length of the tibial condyle (G-I) and medial 1/3 region of the tibia condyle (J-L) for naïve (3-week: n = 8, 6-week: n = 5 - 8, 12-week: n = 3) (blue), sham-operated (3-week: n = 8, 6-week: n = 5 - 6, 12-week: n = 3) (red) and MMT-operated (3-week: n = 8, 6-week: n = 7 - 8, 12-week: n = 3 - 5) (black) hindlimbs. No significant changes to articular cartilage parameters were found between sham and naïve groups and over time. Data points were mean ± SD. b – significance between MMT and sham groups at corresponding timepoint, c - significance between MMT and naïve groups at corresponding timepoint. Black bars above the data represent significant differences between timepoints within the MMT group. All marked significances were *p* < 0.05.

**Figure 5:**
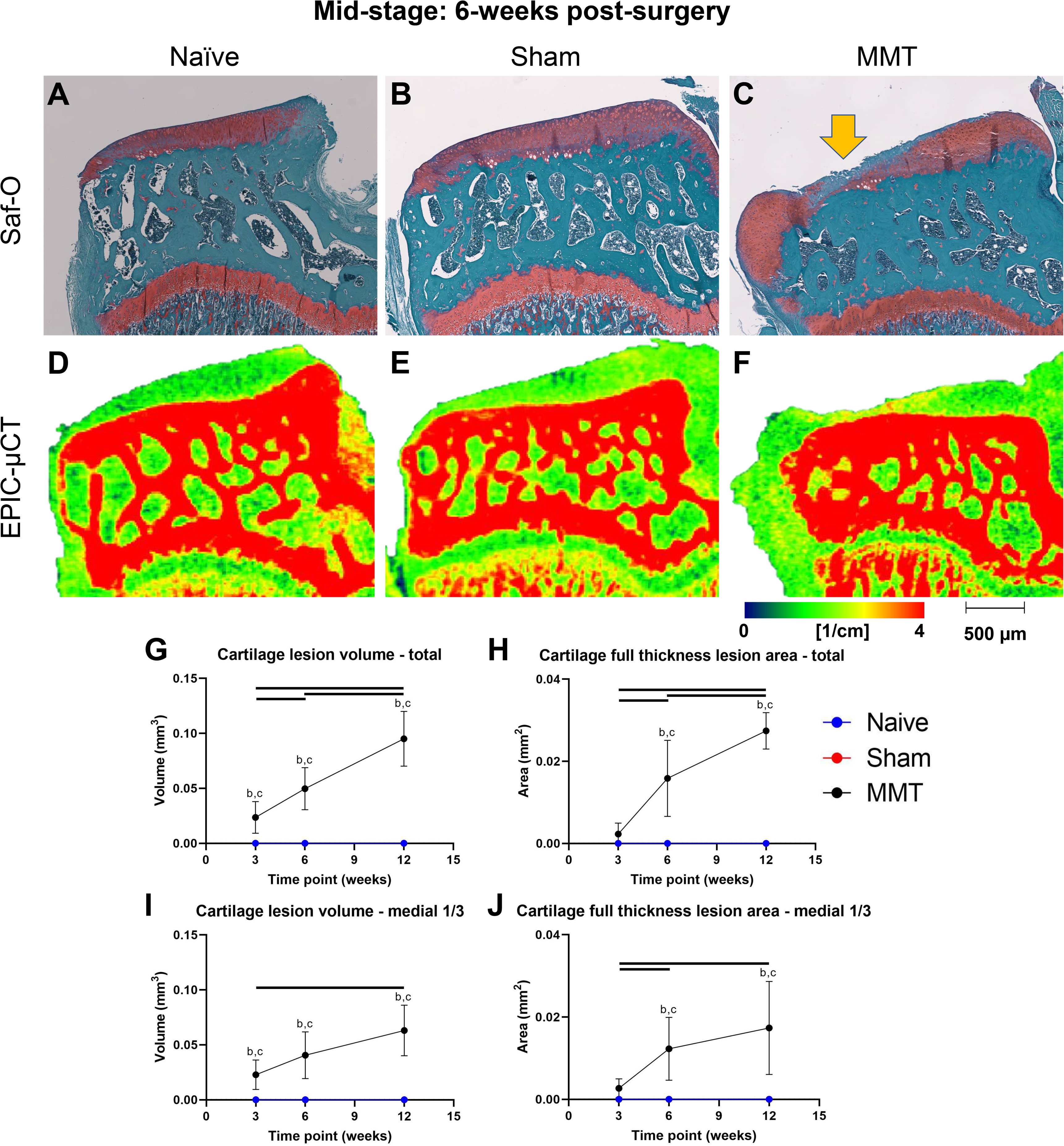
Development of cartilage lesions and exposure of subchondral during OA progression. Rats were euthanized at 6-weeks following Sham- or MMT-surgery and their hindlimbs were fixed in 4% neutral-buffered formalin. After embedding in paraffin, 5μm coronal serial sections of naïve (A), Sham-operated (B) and MMT-operated (C) tibiae were placed onto glass slides and stained with safranin-O/fast-green. Red safranin-O stain was found in the articular cartilage layer and growth plate. These regions were dominated by chondrocytes and the red color was indicative of proteoglycan matrix. Fast-green was used as a counter stain. Lesions (yellow arrow) were evident in the articular cartilage layer of MMT-operated hindlimbs and were not found in naïve or Sham-operated hindlimbs. 2D images of EPIC-μCT coronal sections that closely matched the safranin-O/fast green histology sections were shown for naïve (D), Sham-operated (E) and MMT-operated (F) tibia. Custom-coded MATLAB algorithms were used to quantify cartilage lesions and full thickness lesion area across the total length of the tibial condyle (G-H) and medial 1/3 region of the tibia condyle (I-J) for naïve (3-week: n = 8, 6-week: n = 8, 12-week: n = 3) (blue), sham-operated (3-week: n = 8, 6-week: n = 6, 12-week: n = 3) (red) and MMT-operated (3-week: n = 8, 6-week: n = 6, 12-week: n = 5) (black) hindlimbs. No lesions were measured for naïve and sham tibiae for any time point after surgery. Data points were mean ± SD. b – significance between MMT and sham groups at corresponding timepoint, c - significance between MMT and naïve groups at corresponding timepoint. Black bars above the data represent significant differences between timepoints within the MMT group. All marked significances were *p* < 0.05.

**Figure 6:**
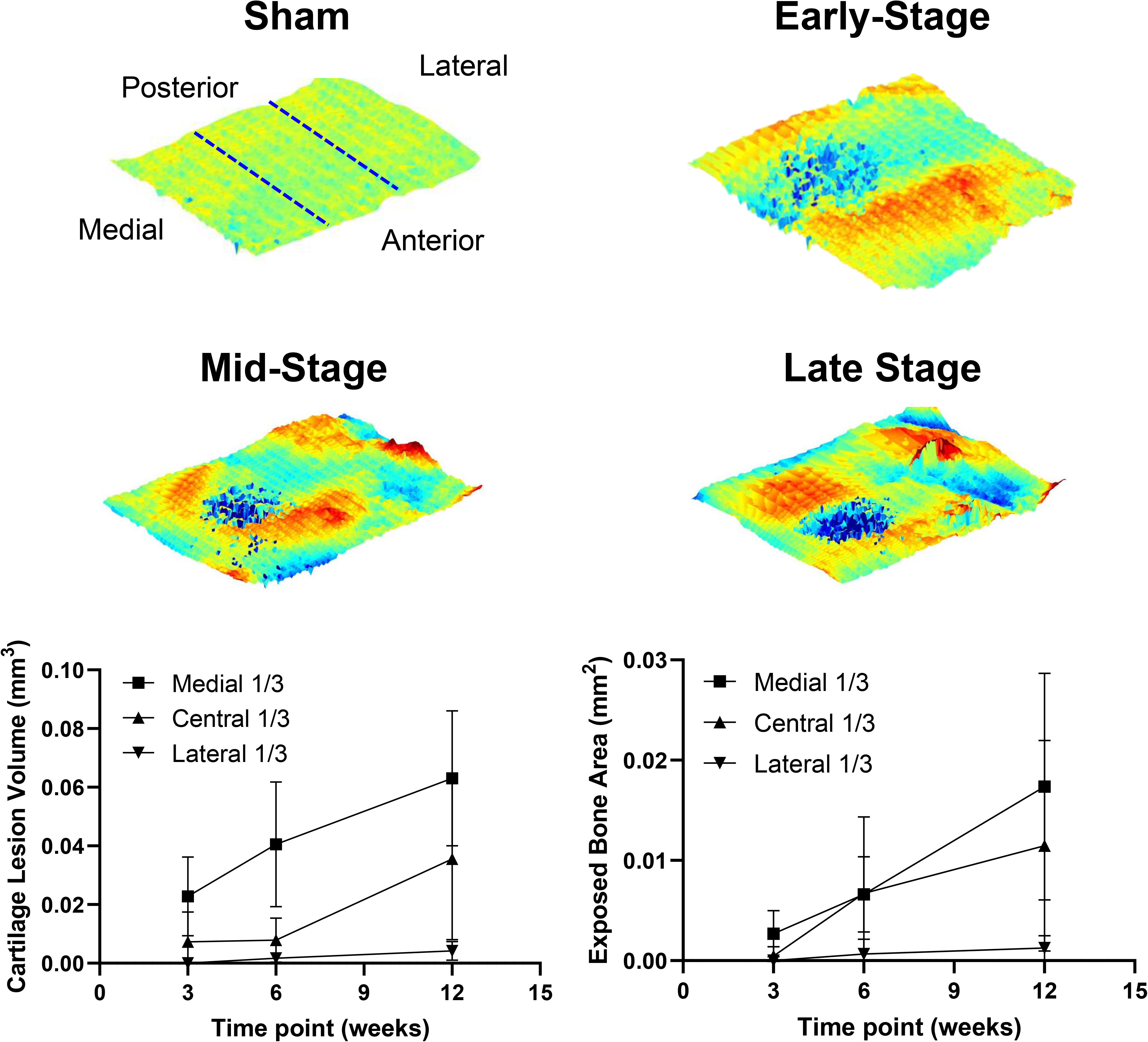
Expansion of lesions from medial to lateral regions of the tibia condyle. 3D rendering of the articular cartilage surface, using our customized MATLAB, illustrated the progression of cartilage lesions and of exposed bone across the tibia condyle for Sham and early-, mid- and late stages of osteoarthritis development. Dark blue represented the area that is devoid of articular cartilage. Custom-coded MATLAB algorithms were used to quantify the cartilage lesions and exposed bone area in the medial, central, and lateral regions of the tibia condyle. Data points were mean ± SD.

Serial slices of the tibiae were stained with hematoxylin and eosin to determine the effect of osteoarthritis on cells within the articular cartilage (Supplementary Figure 1). There were less stained nuclei within the articular cartilage of MMT-operated hindlimb compared to naïve and Sham-operated hindlimbs. Quantification of individual cells was not possible with EPIC-μCT technique and thus, changes to individual cells within the tibiae were not pursued herein.

### Qualitative and quantitative analysis of cartilage lesions and exposed bone

In addition to surface roughness, our custom MATLAB algorithm calculated the lesion volume and full-thickness lesion area (exposed bone) of MMT-operated tibia, which were not evident in any of the naïve or Sham-operated tibiae, even out to 12-weeks post-surgery (Fig. 5G-J). At 6-weeks post-surgery, cartilage lesions (marked by yellow arrow in Figure 5C) were prominent in the tibia of MMT-operated hindlimbs and in some instances, resulted in the loss of chondrocytes down to the subchondral bone layer. The summation of lesion volume across the articular cartilage layer of MMT-operated hindlimbs progressively increased post-surgery. The 3D rendering of our MATLAB algorithm (Figure 6) illustrated the dominance of lesions (dark blue) within the medial 1/3 region and expanded radially towards other regions (i.e., central 1/3 and lateral 1/3) of the condyle at later time points.

The continued burrowing of lesions led to the total loss of chondrocytes within the articular cartilage layer and resulted in exposing the subchondral bone, which was quantified as cartilage full-thickness lesion area. The exposed bone area of MMT-operated hindlimbs increased over time post-surgery and resulted in reduced articular cartilage at 12-weeks post-surgery, especially within the medial 1/3 region (Figure 6).

### Qualitative and quantitative analysis of osteophyte formation

Osteophytes are bone spurs that typically originate along the marginal edges of the proximal tibia during knee osteoarthritis. In the rat MMT model, there was evidence of cartilaginous osteophyte formation preceding mineralization (Figure 7), supporting the notion of endochondral ossification. Using safranin-O to stain for proteoglycan, there was protrusion of chondrocytes along the medial edge of MMT-operated hindlimbs (Figure 7C). The recognition of osteophytes may be subjective from 2D μCT images, thus we used the growth plate as a point of reference of “normal” marginal edge of the tibia (red line Figure 7C). With EPIC-μCT, the cartilaginous osteophytes were distinguishable from mineralized osteophytes (higher attenuation) because of the binding of the contrast agent Hexabrix 320 to the extracellular proteoglycan matrices and was distinguishable from neighboring soft tissue and air (lower attenuation) (Figure 7F). Total osteophytes in MMT-operated hindlimbs were evident at 3-weeks post-surgery and reached a plateau at 6- and 12-weeks. The volume of cartilaginous osteophytes was an order of magnitude higher than mineralized osteophytes of MMT-operated hindlimbs. Further, there was a significant reduction of cartilaginous osteophyte at 12-weeks post-surgery, and this was coincident with a significant increase of mineralized osteophyte. There was a significant difference between Sham and naïve hindlimbs with respect to cartilaginous osteophyte volume; while no significant differences between Sham and naïve were found for any of the previously described articular cartilage parameters, e.g., thickness, volume, surface roughness, lesions, and exposed bone.

**Figure 7:**
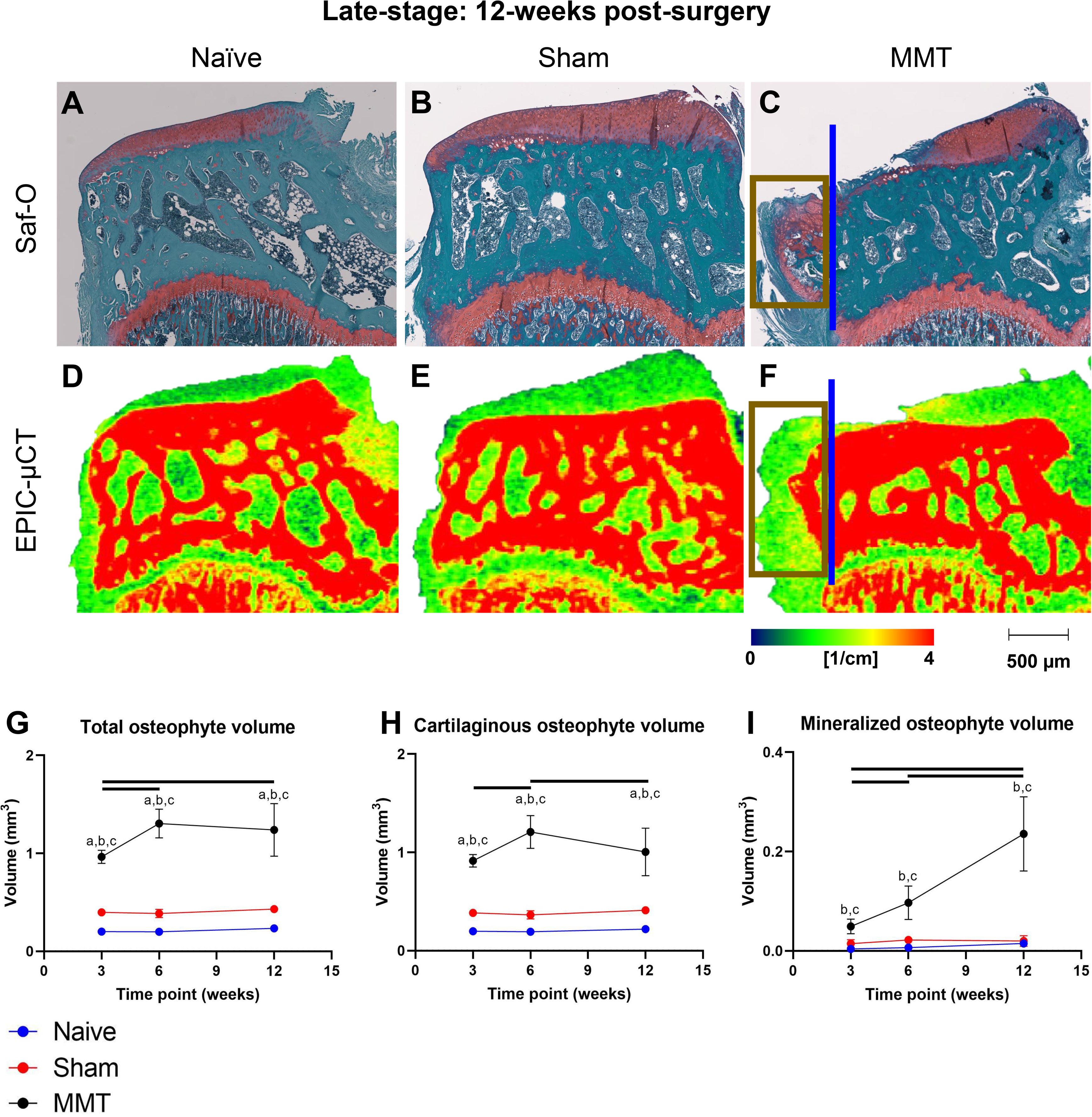
Development of osteophytes during OA progression. Rats were euthanized at 12-weeks following Sham- or MMT-surgery and their hindlimbs were fixed in 4% neutral-buffered formalin. After embedding in paraffin, 5μm coronal serial sections of naïve (A), Sham-operated (B) and MMT-operated (C) tibiae were placed onto glass slides and stained with safranin-O/fast-green. Red safranin-O stain was found in the articular cartilage layer, growth plate and osteophytes. These regions were dominated by chondrocytes and the red color was indicative of proteoglycan matrix. Fast-green was used as a counter stain. Osteophytes (yellow box) were evident along the medial, marginal edge of MMT-operated hindlimbs. The growth plate was used as a point of reference for “normal” marginal edge of tibia (red line). 2D images of EPIC-μCT coronal sections that closely matched the safranin-O/fast green histology sections were shown for naïve (D), Sham-operated (E) and MMT-operated (F) tibia. Custom-coded MATLAB algorithms were used to quantify cartilaginous (H) and mineralized (I) osteophytes and their summation (G, Total) for naïve (3-week: n = 8, 6-week: n = 8, 12-week: n = 3) (blue), sham-operated (3-week: n = 8, 6-week: n = 7, 12-week: n = 3) (red) and MMT-operated (3-week: n = 8, 6-week: n = 8, 12-week: n = 5) (black) hindlimbs. Data points were mean ± SD. a – significance between naïve and sham groups at corresponding timepoint, b – significance between MMT and sham groups at corresponding timepoint, c – significance between MMT and naïve groups at corresponding timepoint. Black bars above the data represent significant differences between timepoints within the MMT group. All marked significances were *p* < 0.05.

### Quantitative analysis of subchondral bone morphometry and structure

EPIC-μCT was used to quantitatively assess subchondral bone thickness and attenuation across the full length and medial 1/3 regions of the medial tibial condyle (Figure 8). Similar to articular cartilage phenotypes, there was no significant difference in subchondral bone measurements between sham and naïve hindlimbs. For MMT-operated hindlimbs, subchondral bone thickness and attenuation gradually increased during the weeks after surgery and were significantly higher than naïve hindlimb at 12-weeks post-surgery.

**Figure 8:**
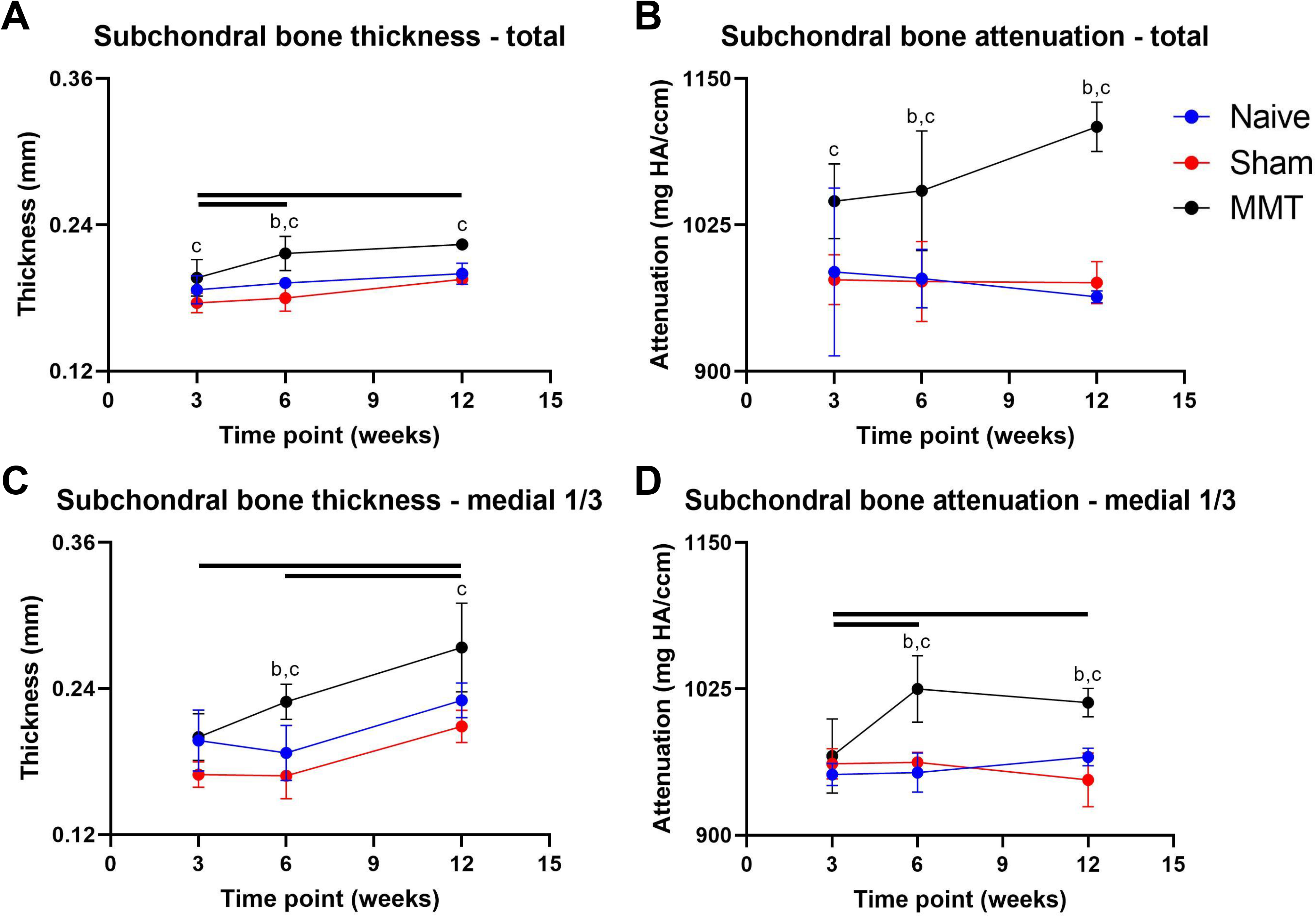
Changes to subchondral bone during OA progression. Custom-coded MATLAB algorithms were used to quantify subchondral bone thickness (A, C) and attenuation (B, D) across the total length of the tibial condyle (A, B) and medial 1/3 region of the tibia condyle (C, D) for naïve (3-week: n = 6 - 8, 6-week: n = 5 - 8, 12-week: n = 3) (blue), sham-operated (3-week: n = 6 - 8, 6-week: n = 4 - 7, 12-week: n = 3) (red) and MMT-operated (3-week: n = 4 - 8, 6-week: n = 5 - 8, 12-week: n = 3 - 5) (black) hindlimbs. No significant changes to subchondral bone parameters were found between sham and naïve groups. Data points were mean ± SD. b – significance between MMT and sham groups at corresponding timepoint, c – significance between MMT and naïve groups at corresponding timepoint. Black bars above the data represent significant differences between timepoints within the MMT group. All marked significances were *p* < 0.05.

## Discussion

The rat MMT model has been extensively used to characterize the onset of post-traumatic osteoarthritis, mostly out to 3-weeks post-surgery and in occasional instances out to 6-weeks post-surgery. Herein, we qualitatively and quantitatively characterized the MMT-induced changes to articular cartilage, subchondral bone, and osteophyte formation at early-(3-weeks post-surgery), mid-(6-weeks post-surgery), and late-stage (12-weeks post-surgery) of osteoarthritis development. At the early stage of osteoarthritis, prominent changes were found in articular cartilage measurements, i.e., thickness, volume, and surface roughness. These changes were quantifiable by using EPIC-μCT, a technique that enables assessment of the articular cartilage. The increased thickness and volume of articular cartilage found in the tibiae of MMT-operated hindlimbs suggested a swelling of the articular cartilage layer, most likely due to the initial degradation of extracellular matrices. From H&E histology, we cannot specifically attribute the increased articular cartilage volume due to expansion or propagation of articular chondrocytes because there was a loss of chondrocytes within the articular layer of MMT-induced osteoarthritis hindlimbs. Further, the thickness and volume of articular cartilage were not significantly different between MMT and Sham (or naïve) hindlimbs at late stage (12-weeks post-surgery) and this was because of the increased loss of articular cartilage at the advanced stage of osteoarthritis in MMT-operated hindlimbs.

Changes to the articular cartilage layer may directly impact the subchondral bone layer, which lies immediately below the articular cartilage. With μCT technique, there was a significant gradual increase in compactness (defined by attenuation) and size (defined by thickness) of the subchondral bone layer of MMT-induced osteoarthritis hindlimbs. These changes were not discernible with standard H&E or safranin-O histology techniques, which was one of the benefits of using μCT for analysis. The increase of subchondral bone in the MMT model was different from the reported decrease of subchondral bone following an anterior collateral ligament tear rat injury model [24–26]. As others have also suggested, the different effects on the subchondral bone may be due to the difference in the post-traumatic osteoarthritis models[27].

Using μCT serial images to quantify key features in osteoarthritis progression, we can compare stages in rat model to key stages in human disease (Figure 9). In particular, the effect of MMT-induction was most prominent in the medial region of the tibia condyle and more specifically, the changes to articular cartilage resembled the degradation of articular cartilage in human osteoarthritis. Histopathology of human osteoarthritis progression was well characterized and described by Mankin and colleagues [28] and later refined by Osteoarthritis Research Society International (OARSI) [29, 30]. We compared the OARSI scoring of human osteoarthritis to findings for rat MMT-induced osteoarthritis disease. Articular cartilage damage is evident within the early stage of rat osteoarthritis and is resembles grades 1-3 of human OARSI scores. The rapid deterioration of articular cartilage in the rat model makes it an ideal model to test potential therapeutics for osteoarthritis. For instance, the use of amniotic membrane alone [13] or in conjunction with umbilical cord tissues [31] may delay osteoarthritis progression when treatment was provided at the early stage of osteoarthritis. Other studies [19, 20] have similarly shown that paracrine secretion from human mesenchymal stromal cells (hMSCs) may delay the onset of osteoarthritis development in the rat MMT model. That study showed that hMSCs increased the size of osteophytes while providing a chondroprotective effect on articular cartilage, suggesting that MSCs may regulate osteoarthritis tissues differently. Taken altogether, the rat MMT-induced osteoarthritis model shares semblance to human osteoarthritis progression. The benefit of the rat osteoarthritis model is its relatively fast onset of osteoarthritis (within first 3 weeks of surgery) and the quantitative method for analysis, making it ideal for testing therapeutics and determining their efficacy in a whole animal.

**Figure 9:**
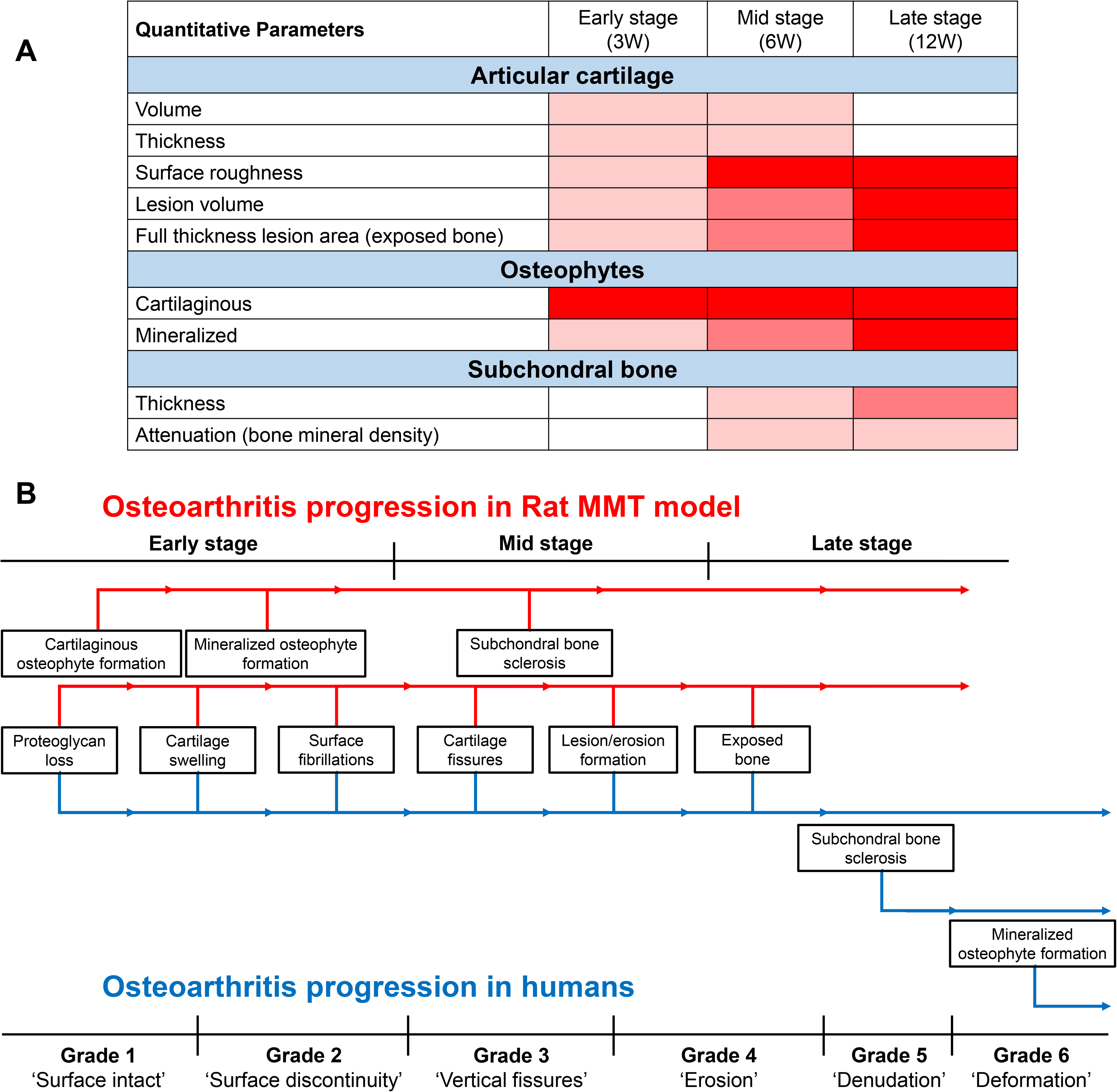
Relation of osteoarthritis progression in rat MMT model and human disease. (A), Summary table for osteoarthritis phenotypes in rat MMT model at early-, mid- and late-stage. The severity of each phenotype is color-coded from minimal (light pink) to gradually being pronounced (red). (B), Comparison of osteoarthritis progression in rat MMT model and human OARSI scores (grade 1 – 6). Damage to articular cartilage is similar for rat MMT model and early changes in human disease pathology. The prominence of osteophyte and changes to subchondral bone occurred earlier in the rat MMT model compared to findings for human osteoarthritis.

Many therapies have looked promising and showed therapeutic benefits in the rat MMT model; however, no therapeutics, to date, have effectively translated into clinical practice. Our study herein shows 3-weeks of MMT-induction in the rat results in early signs of osteoarthritis, which is unlikely a stage when symptomatic osteoarthritis patient seeks and receives treatment. Instead, MMT-induction out to 12-weeks post-surgery shows osteoarthritis deterioration closer to OARSI score of 4-6, which may be the point when osteoarthritis patient may exhibit pain and eventual immobility. As therapeutics are further developed, treatment efficacy should be initiated at mid- and/or late- stage of the rat MMT animal model and this would be a closer match to clinical relevance.

## Acknowledgement

The authors thank Mila Friedman for providing the histology samples.

---

**Supplementary Figure 1:**
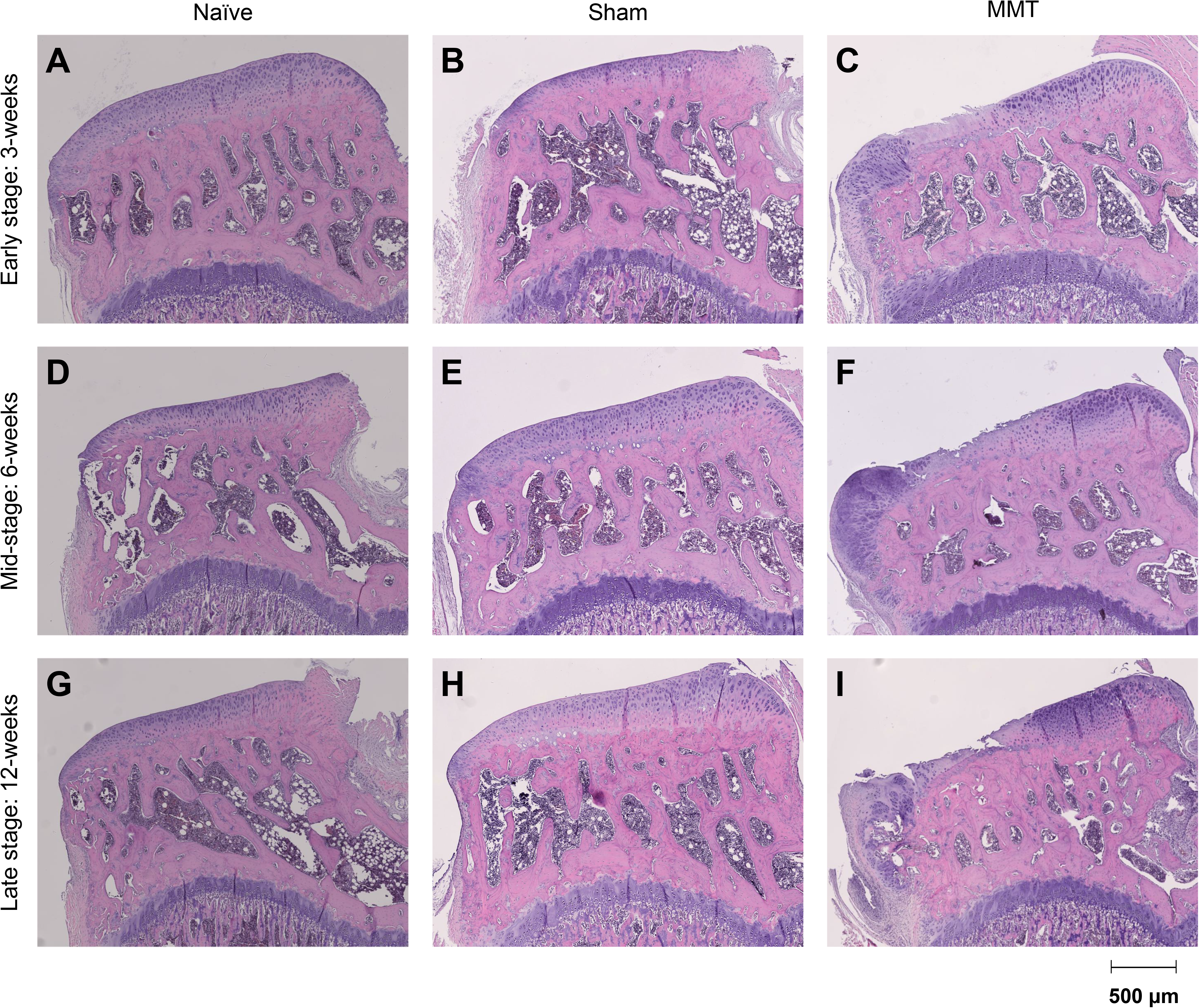
Loss of chondrocytes within the articular cartilage layer during OA progression. Rats were euthanized at 3-, 6- and 12-weeks following Sham- or MMT-surgery and their hindlimbs were fixed in 4% neutral-buffered formalin. After embedding in paraffin, 5μm coronal serial sections of naïve (A), Sham-operated (B) and MMT-operated (C) tibiae were placed onto glass slides and stained with hematoxylin and eosin (H&E). The representative H&E images were the adjacent serial section of the safranin-O/fast green images found in Figures 4, 5 and 6. Hematoxylin stained the nuclei of cells and eosin stained the cytoplasm of cells and the extracellular matrices.

